# Fluorescent anionic cyanine plasma membrane probes for live cell and *in vivo* imaging

**DOI:** 10.1101/2024.10.17.618786

**Authors:** Dmytro I. Danylchuk, Igor Khalin, Yelisetty V. Suseela, Severin Filser, Nikolaus Plesnila, Andrey S. Klymchenko

## Abstract

Proper staining of cell plasma membrane is indispensable for fluorescence imaging. Herein, we present an array of five anionic cyanine-based turn-on plasma membrane probes with emission spanning from green to near infrared. They are analogous of commonly used MemBright probes family, where two zwitterionic anchor groups are replaced with anionic sulfonates with dodecyl chains. The developed probes provide selective wash-free staining of plasma membranes of live cells in vitro, featuring improved brightness and slower internalization inside the cells. In comparison to protein-based (wheat germ agglutinin) membrane markers, new membrane probes provide better staining in poorly assessable cell-cell contacts. A key challenge is to stain cell plasma membranes directly in vivo. During *in vivo* brain tissue imaging in living mice by two-photon microscopy, the anionic cyanine probes allowed us to visualize in detail the pyramidal neurons with high image quality, clearly resolving neuron soma, dendrites with dendritic spines and axons with axonal boutons. The developed anionic cyanine-based plasma membrane probes constitute an important extension of the toolbox of fluorescent probes for plasma membrane research.

## Introduction

Plasma membrane (PM) plays an important role in a number of essential biological processes including cell trafficking and migration, signaling, cellular uptake, neural communication and muscle contraction, alongside with its main function of delimiting and separating cell interior.^1^

Visualization of cell membrane is of an increased importance in bioimaging, as it allows to identify and delimit the cells. Moreover, the morphology of PM can provide the information regarding various biological processes such as cell division or death,^2^ also indicating cell damage.^3^ Therefore, imaging of plasma membrane is important for both biological research and early medical diagnosis.^4^ Fluorescence imaging is also a powerful tool for neuroscience,^5^ with the visualization of PM being of particular importance due to its capability to elucidate the details of neuronal organization and membrane-related functioning.^6^ Therefore, development of new fluorescent probes for biomembranes is a rapidly expanding research direction.^7^ Ideal plasma membrane probe must meet multiple criteria in order to be accepted by a broad community of biologists, which includes (1) simplicity of use; (2) high brightness and photostability; (3) specificity of labelling; (4) persistent PM staining with minimal internalization. Additional challenges stem from applications of membrane probes *in vivo*, because of complexity of extracellular medium in tissues, probe elimination, as well as poor tissue penetration by light.

Several approaches towards staining PM have been developed, including genetically encoded proteins and molecular probes. The main advantages of the latter include facile application, alleviating the need to perform genetic modification of cells prior to imaging, together with high brightness and photostability, which can be achieved when using organic fluorophores.^8^ The main challenge in creation of molecular probes for PM is to obtain specific and persistent staining.

Fluorescently labeled lectins, notably wheat germ agglutinin (WGA) and concanavalin A, are commonly used fluorescent membrane probes with high efficiency. However, the main disadvantage of these probes is significantly larger probe size when compared to molecular probes. Moreover, WGA binds N-acetyl-D-glucosamine and sialic acid,^9^ and the direct positioning of fluorophore inside plasma membrane is not assured, which might affect the imaging of plasma membrane lipids. The small size of fluorescent probes with chemical targeting groups coupled with high lipid bilayer affinity allows their precise localization in the lipid bilayer, which improves the quality of PM visualization and is indispensable for studies of the lateral lipid organization in biomembranes.^10^

Highly hydrophobic cyanine dyes, such as DiI, DiO or DiD and PKH family, which are widely used as efficient markers in membrane models, such as liposomes, generally exhibit poor performance when staining live cell membrane due to aggregation and precipitation before reaching the membrane and facile crossing of the PM, leading to indiscriminate staining of all lipid compartments in the cell.^10a,11^ Therefore, specific PM staining requires the use of specially designed targeting anchor groups.^7c^ Generally, chemical PM-targeting groups comprise of a hydrophobic fragment, being either an alkyl chain or the dye itself, and a cationic^12^ or zwitterionic^13^ charged part. On the other side, introduction of negative charges generally increases dye solubility and leads to improved imaging performance.^14^ Recently, we have found that utilization of negatively charged targeting group for a Nile Red probe results in an increased membrane affinity and reduced internalization^15^ compared to its zwitterionic counterpart.

The probes for PM were created using various fluorophores, including BODIPY^12e,12f,13e^ and BF_2_-azadipyrromethene,^14c^ Nile Red,^13c,15^ Prodan,^16^ chromone,^13a,13b^ coumarin,^17^ push-pull fluorene,^18^ phenyl fluorene,^19^ styrylpyridinium,^12a,12c,12d,20^ oligothiophene,^14b,21^ perylene,^22^ fluorescein,^23^ isoindoledione,^24^ purine,^12b^ a variety of aggregation-induced emission dyes,^12b,25^ conjugated polymers,^26^ squaraine^27^ and cyanines.^3,12a,13d^.

Cyanine dyes are highly popular fluorophores for bioimaging applications^8,28^ due to their wide spectral range covered with emission from green to near infrared, relatively high brightness and photostability. Additionally, cyanines are able to form non-fluorescent J-aggregates, which provides an opportunity to create fluorogenic probes.^29^ Utilization of special blinking buffers induces blinking behavior in cyanine dyes,^30^ which, in turn, allows to use them in super-resolution microscopy.^31^ Cyanine dyes are also compatible with two-photon excitation microscopy (2PM), which makes them promising candidates for tissue and *in vivo* imaging.^13d,32^ In our previous work, a series of six cyanine PM probes with zwitterionic targeting groups was synthesized, covering the spectral range from orange to near infrared.^13d^ In addition to bright and photostable labeling in conventional microscopy, these probes were also used for super-resolved imaging of the neck of dendritic spines.^13d^ Moreover, MemBright probe labeled neurons in a brighter manner compared to surrounding cells, resulting in identification of neurons in acute brain tissue sections and neuromuscular junctions.^13d^ However, their application in vivo for labelling cell PM, including neurons has not been realized so far. Recently, PK Mem dyes, analogues of MemBright probes with triplet quencher moiety for minimizing probe phototoxicity, were reported and successfully applied for in vitro and in vivo imaging, including mice brain.^33^ It is important to note that, the zwitterionic targeting motifs do not affect the overall probe charge, and as the cyanine part bears an intrinsic positive charge, the overall probe charge of MemBright and PK Mem remains positive. In principle, the net positive charge could lead to some issues, such as increased dye internalization, non-specific binding to surfaces (both glass and plastic) and rapid elimination in vivo. We hypothesized that the efficiency of PM staining in vitro and in vivo by cyanine dyes could be potentially increased through the utilization of negatively charged targeting groups.

In the present work, we report an array of anionic cyanine-based plasma membrane probes for live cell and *in vivo* imaging.

## Results and discussion

An array of anionic cyanine probes was synthesized (Fig. 1A) with emission spanning from green to NIR. Cyanine dye based benzoxazole heterocycle and short (C3) polymethine bridge (analogue of DiO, Cy2) was chosen as a green emitting fluorophore. The analogues with 1H-indole moieties (analogue of DiI, Cy3) and 1H-benz[e]indole (Cy3.5) heterocycles were chosen to operate in orange and red emission range, respectively. Finally, cyanines with longer polymethine bridge length (C5) were also selected to shift dye emission to far red (Cy5, analogue of DiD) and NIR (Cy5.5) emission. To target these dyes to plasma membranes, two 3-(dodecylammonio)propane-1-sulfonate moieties were chosen, because of their negative charge and proven targeting efficacy.^15-16^ The probes were synthesized in five steps through the corresponding cyanine diacids (Fig. 1A, Scheme S1). These probes are close analogues of MemBright family of dyes, reported by us before. The key difference is the use of negatively charged anchors, which provide the new probes net negative charge (−1) vs net positive charge (+1) of the original MemBright family.^13d^

**Fig. 1.**
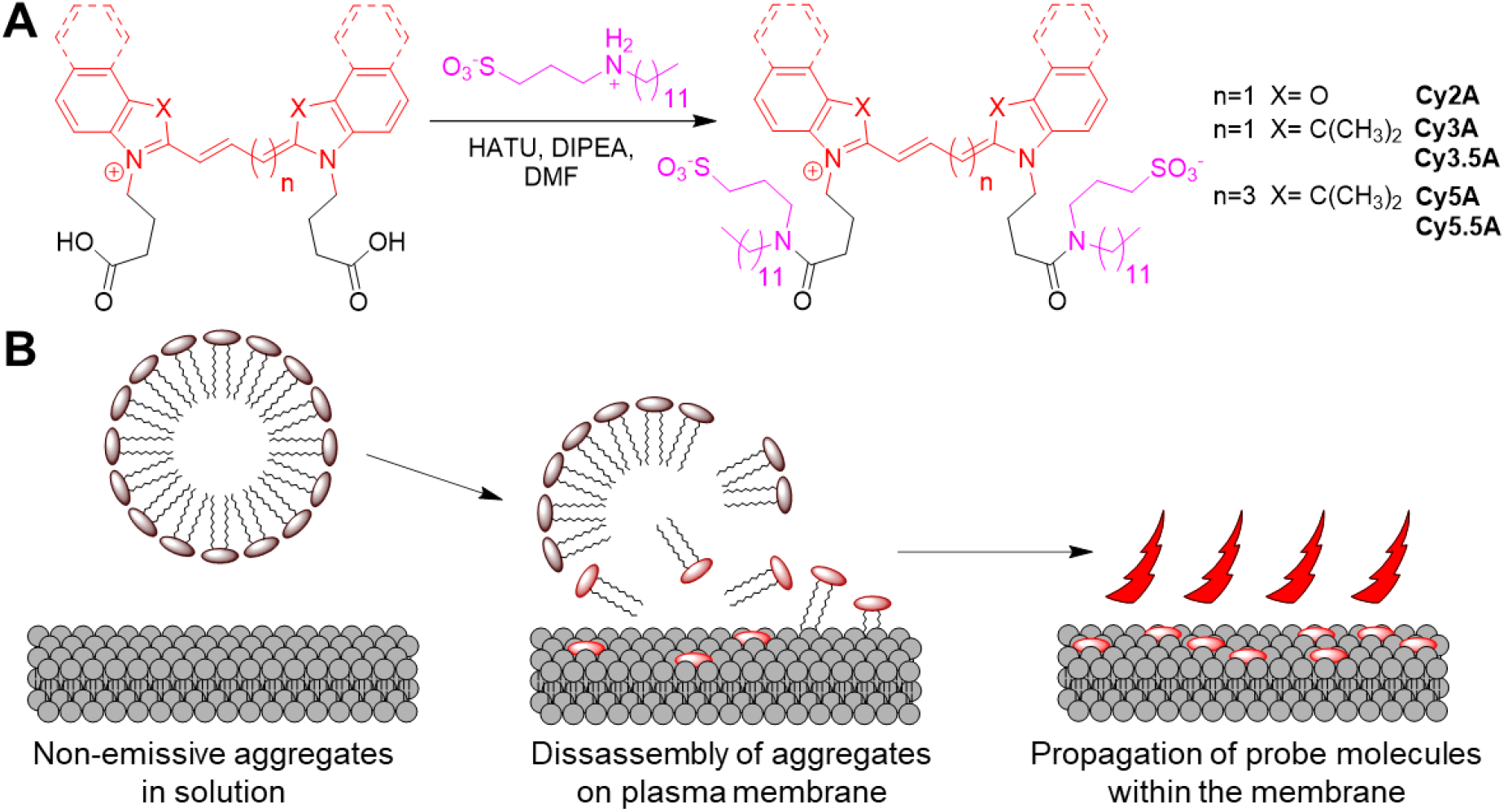
Synthesis (A) and fluorescence turn-on mechanism (B) of anionic cyanine probes.

Due to the amphiphilic nature of the targeting groups the anionic cyanine probes are expected to form non-emissive aggregates in water, which will disassemble in presence of lipid membranes, thus providing a fluorogenic turn-on response (Fig. 1B), similarly to the previously developed MemBright probes (Fig. S1).^13d^

Spectroscopy experiments with anionic cyanine probes were performed in organic solvents and large unilamellar vesicles (LUVs) (Fig. 2, Fig S2-S3). The absorption spectra of anionic cyanine probes in organic solvents (Fig. S2) and DOPC LUVs (Fig. 2A) were narrow and close to the mirror image of their emission spectra (Fig. 2B and Fig. S3), indicating no sign of fluorophore perturbation or aggregation. On the contrary, the probes in phosphate buffer (PB) exhibited an increase in a short-wavelength shoulder in absorption spectra (Fig. 2A, Fig. S2) accompanied by a significant decrease in fluorescence intensity (Fig. 2B, Fig. S3) suggesting the formation of H-aggregates^34^ in aqueous media.

**Fig. 2.**
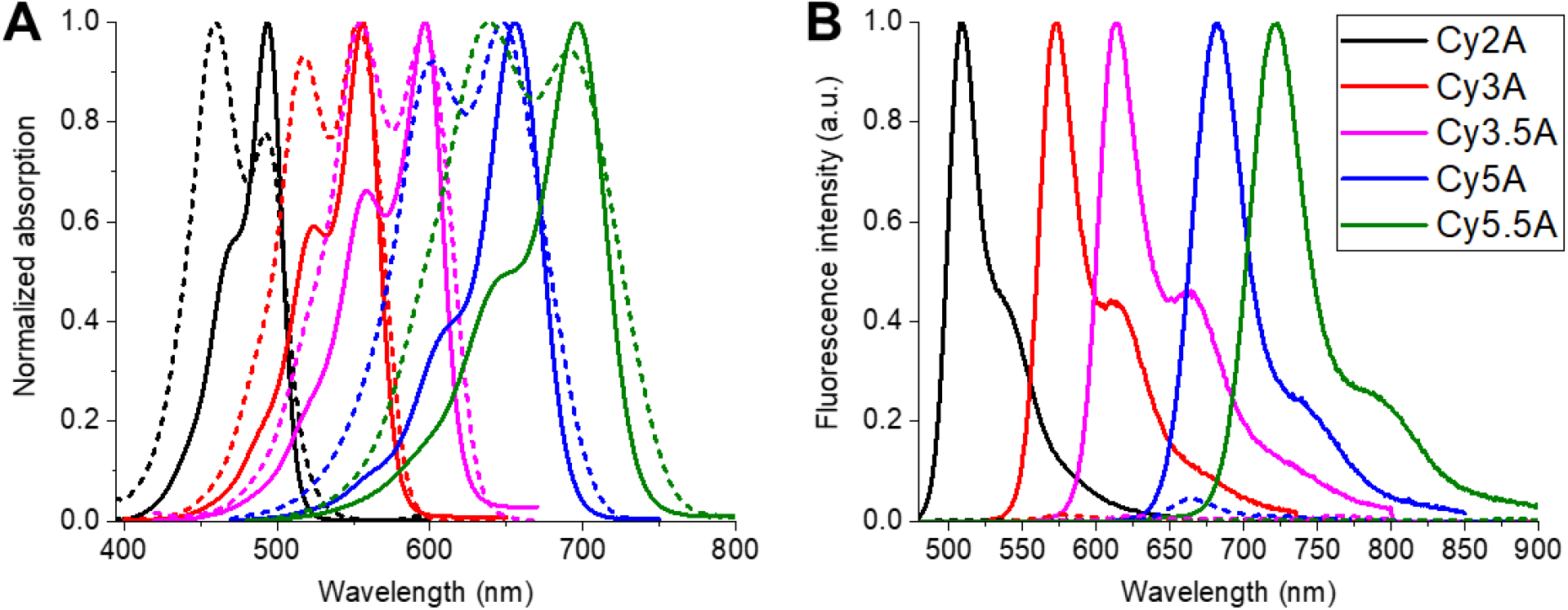
Normalized absorption (A) and emission (B) spectra of anionic cyanine probes. Solid lines represent the spectra in DOPC, while the corresponding spectra in phosphate buffer are represented in dashed lines. Probe concentration was 1 µM for Cy2A and Cy3 and 0.5 µM for Cy3.5, Cy5 and Cy5.5. Total lipid concentration was 400 µM. Excitation wavelength was 470 nm for Cy2A, 540 nm for Cy3A, 546 nm for Cy3.5A, 601 nm for Cy5A and 623 nm for Cy5.5A.

Anionic cyanine probes exhibit relatively high quantum yields (QY) in lipid vesicles and organic solvents (Table 1), being 40 to 190-fold higher for the vesicles compared to those in phosphate buffer. Low QY values in PB (Table 1) further evidence probe aggregation in aqueous medium. Thus, our probes provide a fluorogenic response to the presence of lipid membranes.

**Table 1.** Spectroscopic properties of anionic cyanine probes in solvents and lipid vesicles.

| Probe | Solvent or medium |  |  |  |  |  |  |  |  |  |  |  |
| --- | --- | --- | --- | --- | --- | --- | --- | --- | --- | --- | --- | --- |
|  | MeOH |  |  | PB |  |  | DOPC |  |  | 1,4-Dioxane |  |  |
| | $\lambda_{\max}$<br>Abs | $\lambda_{\max}$<br>Em | QY,<br>% | $\lambda_{\max}$<br>Abs | $\lambda_{\max}$<br>Em | QY | $\lambda_{\max}$<br>Abs | $\lambda_{\max}$<br>Em | QY,<br>% | $\lambda_{\max}$<br>Abs | $\lambda_{\max}$<br>Em | QY,<br>% |
| Cy2A | 486 | 502 | 10.4 | 460 | 697 | 0.9 | 494 | 509 | 47.0 | 495 | 509 | 36.0 |
| Cy3A | 550 | 564 | 11.0 | 553 | 577 | 0.4 | 556 | 573 | 37.2 | 556 | 570 | 34.5 |
| Cy3.5A | 589 | 608 | 24.3 | 555 | 763 | 1.0 | 597 | 614 | 64.4 | 594 | 609 | 52.8 |
| Cy5A | 645 | 669 | 42.8 | 648 | 663 | 1.5 | 656 | 682 | 58.2 | 660 | 682 | 56.4 |
| Cy5.5A | 682 | 707 | 22.4 | 639 | 890 | 0.1 | 696 | 723 | 19.2 | 694 | 721 | 21.6 |

Probe concentration was 1 $\mu\text{M}$ for Cy2A and Cy3 and 0.5 $\mu\text{M}$ for Cy3.5, Cy5 and Cy5.5. Total lipid concentration was 400 $\mu\text{M}$ . Excitation wavelength was 470 nm for Cy2A, 540 nm for Cy3A, 546 nm for Cy3.5A, 601 nm for Cy5A and 623 nm for Cy5.5A.

Next, the new probes were studied in live breast cancer cells MDA-MB-231 by fluorescence microscopy in the presence of reference membrane marker WGA-Alexa 488 conjugate (Fig. 3). We found that the new probes display high selectivity towards plasma membrane, being well co-localized with the reference marker. Cy2A, which could not be used with WGA-Alexa 488, was localized with Cy5A, confirming its PM selectivity. Careful inspection of the images of new dyes vs WGA-Alexa 488 revealed that in the cell-cell contacts the fluorescence intensity of WGA-Alexa 488 was poorly visible, in contrast to staining by all new cyanine probes. Therefore, in the composite images, the cell-cell contacts appeared in red. We assume that in the contacts the narrow space between cells restricts free diffusion of WGA-Alexa 488 and further access of plasma membranes in these areas. Moreover, WGA-Alexa 488 already bound to glycoproteins (to N-acetyl-D-glucosamine and sialic acid residues^9^) probably displays restricted diffusion in these contacts, thus explaining the lack of staining. In contrast, cyanine dyes, which are bound to lipids of plasma membrane, can freely diffuse within the biomembrane and readily access these hidden regions of plasma membranes. Thus, homogenous cell staining even within tightly bound cells constitutes an important advantage of the new probes compared to WGA-Alexa 488. Similar experiments were conducted on KB cells, showing good colocalization of the new cyanine probes with WGA-Alexa 488 (Fig. S4). In this case, no differences were observed in cell-cell contacts, probably because they were not as tight as for MDA-MB-231 cells. One could also notice staining with WGA-Alexa 488 was homogenous compared to new cyanine dyes, so that some variations in relative fluorescence intensities in PM of the different cells in the merged images was observed. It could arise from the differences in the PM binding mechanism of WGA vs cyanine probes, together with the significantly larger probe size of WGA conjugates when compared to cyanine probes.

**Fig. 3.**
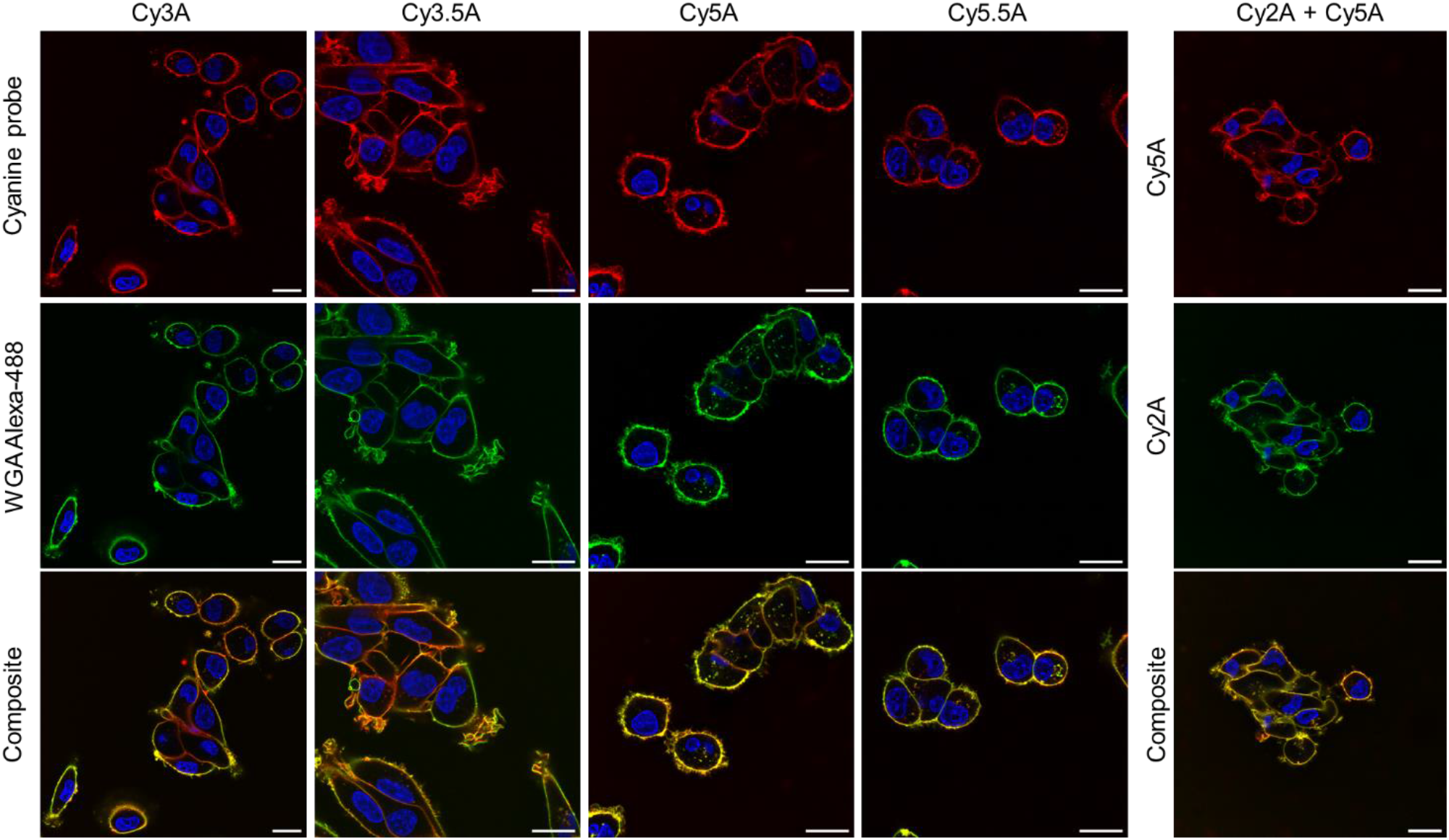
Colocalization of anionic cyanine probes (50 nM; red) with WGA Alexa 488 conjugate (50 nM; green) in live MDA-MB-231 cells incubated for 10 min at room temperature. Hoechst 33342 (1 μM; blue) was used as a nuclear staining probe. Scale bar set to 20 μm.

We also made a direct comparison of new cyanine probes Cy3A and Cy5A with corresponding MemBright probes MB-Cy3 and MB-Cy5 after incubation for 15 min at RT with MDA-MB-231 cells (Fig. 4). First, both new probes showed higher brightness on cell plasma membranes compared to their MemBright analogues. Second, in these conditions, both MB-Cy3 and MB-Cy5 showed some intracellular dots, which according to our previous studies could be assigned to endosomes.^13d^ In contrast, new probes showed practically no sign of internalization after this incubation period. These results corroborate with our observations with WGA-Alexa 488 and previous data, confirming that the anionic probe Cy3A internalizes slower than cationic MB-Cy3. Our previous work showed that MB-Cy3 and MB-Cy5 internalized slightly faster than WGA-Alexa 488,^13d^ while the new probes showed practically the same slow internalization as WGA-Alexa 488 after short incubation (see Fig. 3). The differences are probably related to the net negative charge of new cyanine probes compared to the net positive charge of MemBright probe family. Thus, the new probes present advantages over both protein-based (WGA) and small molecule membrane probes.

**Fig. 4.**
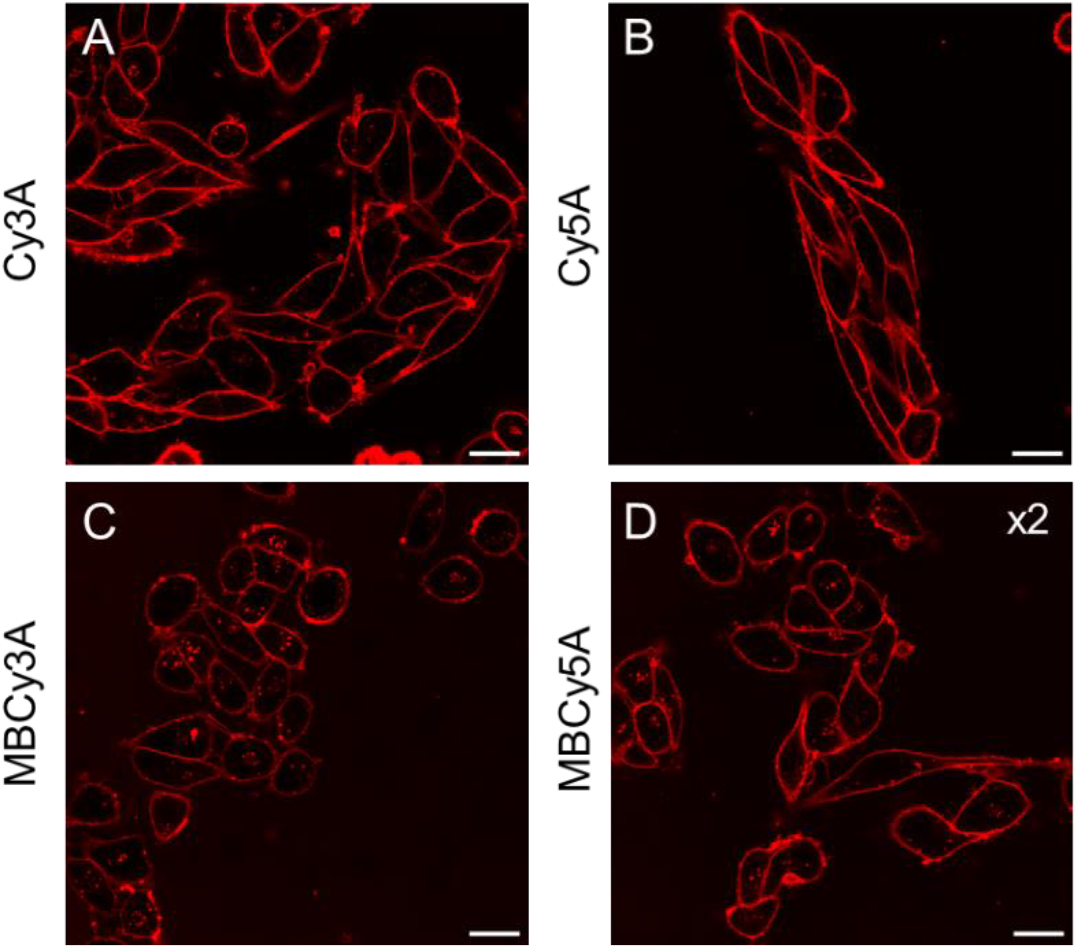
Confocal microscopy images of live MDA-MB-231 cells treated with Cy3A (A), Cy5A (B), MemBright-Cy3 (C) and MemBright-Cy5 (D) for 15 min at room temperature. The concentration of all probes used was 25 nM. Scale bar set to 20 μm.

Fluorescence imaging of plasma membranes *in vivo* poses additional challenges due to a complex and heterogeneous environment, including cells, extracellular matrix and biological fluids together with non-negligible sample thickness, leading to increased autofluorescence and increased absorption and scattering of light.^3,32a,35^ Two-photon excitation microscopy (2PM) minimizes photobleaching, phototoxicity, and cell autofluorescence while also providing 3D sectioning by concentrating excitation at a near-infrared laser’s focal point.^36^ In addition, 2PM significantly reduces light scattering, enhancing tissue penetration depth, crucial for live brain imaging.^37^ Here, we challenged our probes in ambitious application for *in vivo* imaging of neurons directly in live mouse.

Previous results with zwitterionic targeting groups (MemBright family) have shown that the probes stained neurons brighter compared to other surrounding cells,^13d^ although *in vivo* applications could not be shown. Cy3.5A was chosen as a promising candidate for *in vivo* brain tissue imaging experiments due to the high two-photon absorption cross-section and red-shifted emission of Cy3.5 fluorophore.^13d^ First, we searched for a delivery agent of Cy3.5A, which would partially decrease dye aggregation in water, leading to a faster dye diffusion form the injection site and rapid staining of neuronal membranes in a larger volume. The solubilization of anionic cyanines in water was monitored by absorption spectra (Fig. S5), where the efficient delivery agents induced a decrease in the short-wavelength shoulder. Addition of Me-β-cyclodextrin also led to a hypsochromic shift of absorption maximum while BSA induced a bathochromic shift; in both cases the absorption bands became more narrow, closer to ones of cyanine dyes solubilized in organic solvents (Fig. S2). Thus, both BSA and Me-β-cyclodextrin were able to solubilize Cy3A, although for the *in vivo* studies we have chosen the latter (simpler) agent.

Next, Cy3.5A probe was applied for *in vivo* 2PM microscopy of layer II-III pyramidal neurons in mouse brain (Fig. 5, S6). When Cy3.5 was directly injected without helping agent, the staining was observed but it was not so homogenous, whereas the formulation with BSA showed rather fuzzy staining (Fig. S6). On the other hand, a solution of anionic cyanine together with Me-β-cyclodextrin was delivered into the mouse cortex through a stereotactic injection (Fig. 5A), and the neurons were imaged through an acute cranial window (Fig. 5B) 30 min post-injection. Cy3.5A provided bright and specific staining of layer II-III pyramidal neurons at 100 µm depth from the brain surface (Fig. 5C). The high quality of the obtained images allowed to distinguish clearly the neuron soma (Fig. 5D), dendrites with dendritic spines (Fig. 5E) and axons with axonal boutons (Fig. 5F). Moreover, the dye showed no sign of photobleaching during these imaging experiments and did not eliminate from the neuronal membrane after one hour of imaging, thus exhibiting a photostable and robust staining.

**Fig. 5.**
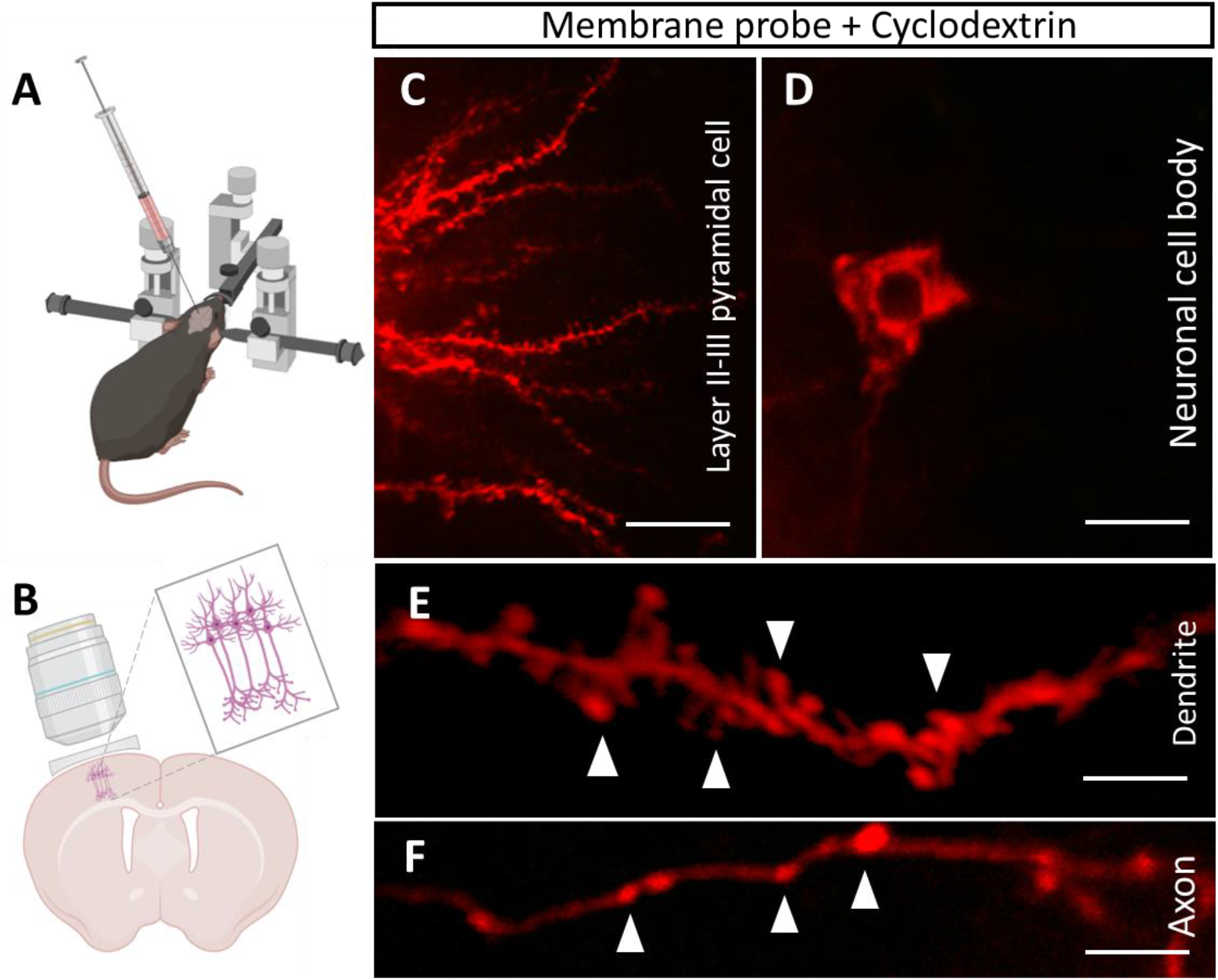
*In vivo* 2-photon microscopy of neurons stained by Cy3.5A probe injected into the brain parenchyma. (A) Stereotactic injection of the probe into the mouse cortex. (B) Intravital imaging of the stained neuronal structures through an acute cranial window. Zoomed insert shows layer II-III pyramidal neurons. (C) Representative micrograph of stained neurites *in vivo* at 100 µm depth from the brain surface. (D) Representative micrograph of stained pyramidal neuron soma. (E) Representative micrograph of stained single dendrite. White arrows indicate dendritic spines. (F) Representative micrograph of stained single axon. White arrows indicate axonal boutons. Scale bar: (C) – 20 µm; (D) – 10 µm; (E, F) – 5 µm.

## Conclusions

In this work, we developed an array of five anionic cyanine probes with the emission spanning from green to near infrared and demonstrated their efficiency for cellular and *in vivo* microcopy. Modification of cyanines with two anionic plasma membrane-targeting motifs allowed to obtain negatively charged probes and ensure selective and persistent staining of the PM. In aqueous media, the probes form non-fluorescent aggregates, which dissociate in presence of lipid membranes, forming highly fluorescent molecular species solubilized in lipid bilayer. This fluorogenic response allowed microscopy imaging in live cells with high signal to noise ratio without any washing step. In comparison to reference membranes marker based on labelled WGA, anionic cyanine probes showed significantly slower internalization. Comparison with a MemBright probe (MB-Cy3) showed that the anionic analogue presents multiple advantages: higher brightness and also slower internalization inside the cells. Thus, the use of anionic anchors, which bring net negative charge to cyanine could improve staining properties compared to the analogues with net positive charge.

Particularly challenging is imaging PM of cells directly in live animals, in particular, for brain neurons. *In vivo* techniques for brain tissue imaging primarily involve the use of Thy-1-GFP mice, producing genetically encoded fluorescent proteins in neurons.^38^ While this approach allows to specifically label neurons and is generally compatible with *in vivo* imaging, the obtained staining is heterogeneous. Moreover, this technique is almost completely reserved for the mouse species since transgenic marmoset, rat, or other mammal animal models are still rare.^39^ The alternative methods used to label neurons in brain tissue include biocytin injection, followed by treatment with fluorescent avidin conjugates,^40^ staining with DiI, DiO or DiD dyes in crystals^41^ and immunolabeling.^41b^ These techniques do not require any use of transfection or transgenic animals, however they are not always compatible with *in vivo* imaging and their specificity to plasma membranes is poor. Additional complication for labeling with long-chain these hydrophobic cyanine probes arises from their poor solubility in water.^38d^ Our results show that the new probes (in particular Cy3.5A) could address at least in part the mentioned issues. During *in vivo* 2PM imaging experiments Cy3.5A probe provided bright, specific and persistent staining of layer II-III pyramidal neurons, resulting in high-quality images with clearly resolved soma, dendrites with dendritic spines and axons with axonal boutons. To the best of our knowledge, this is the first example of neuronal membranes imaged at this level of detail in live mouse brain using an exogenous molecular fluorescent probe. Recent attempt to do so resulted in neuron images, but without details on their dendrite structure.^33^ On the other hand, an important advantage of our probes over fluorescent proteins is that they can in principle be applied to any sample, not being restricted to transgenic mouse species. Moreover, due to local staining protocol, these probes could provide labelling of specific neurons in brain.

Overall, due to availability of five different colors within a spectral range of 500-750 nm combined with high brightness, selectivity and persistence of staining, high signal-to-noise ratio and compatibility with cellular and *in vivo* imaging, anionic cyanine probes have a great potential as a powerful toolkit for plasma membrane imaging in cell biology and neuroscience.

## Acknowledgements

This work was supported by the European Research Council ERC Consolidator grant BrightSens 648528 and ED222 of University of Strasbourg and Deutsche Forschungsgemeinschaft (DFG, German Research Foundation) – projects number: 457586042. Philippe Chabert is acknowledged for help with synthesis of some starting cyanine derivatives.

## Author Information

### Author Contributions

The manuscript was written through contributions of all authors. All authors have given approval to the final version of the manuscript.

### Notes

The authors declare no competing financial interest.

Supporting information available: Materials and Methods section and additional supporting figures.

