## Supporting Information for "Fluorescent anionic cyanine plasma membrane probes for live cell and *in vivo* imaging"

### Materials and methods

**Materials and characterization of compounds.** All the reagents were purchased from Sigma-Aldrich, Alfa Aesar, TCI or Thermo Fisher Scientific and used as received. MilliQ-water (Millipore) was used in all experiments. NMR spectra were recorded at 20°C on a BrukerAvance III 400 MHz spectrometer. Mass spectra were obtained using an Agilent Q-TOF 6520 mass spectrometer. Absorption and emission spectra were recorded on an Edinburgh FS5 spectrofluorometer equipped with a thermostated cell holder. Fluorescence quantum yields were measured using: Fluorescein in 0.1 M NaOH ($\lambda_{ex}$= 470 nm, $QY_{ref}$ = 0.91)[1] ; Rhodamine 6G in EtOH ($\lambda_{ex}$= 510 nm, $QY_{ref}$ = 0.94)[2]; Cresyl violet in MeOH ($\lambda_{ex}$= 546 nm, $QY_{ref}$ = 0.65)[3]; Cresyl violet in EtOH ($\lambda_{ex}$= 601 nm, $QY_{ref}$ = 0.67)[3]; Rhodamine800 in EtOH ($\lambda_{ex}$= 623 nm, $QY_{ref}$ = 0.25)[4] as a reference for Cy2A, Cy3A, Cy3.5A, Cy5A and Cy5.5A, correspondingly.

**Synthesis.** The synthesis of anionic cyanines was performed starting from either 2-methybenzoxazole or 1,1,2-Trimethyl-1H-benzo[e]indole (Fig. S1). Compounds **3a**,[5] **3b**[5] and **2d**[6] were synthesized according to the literature procedures.

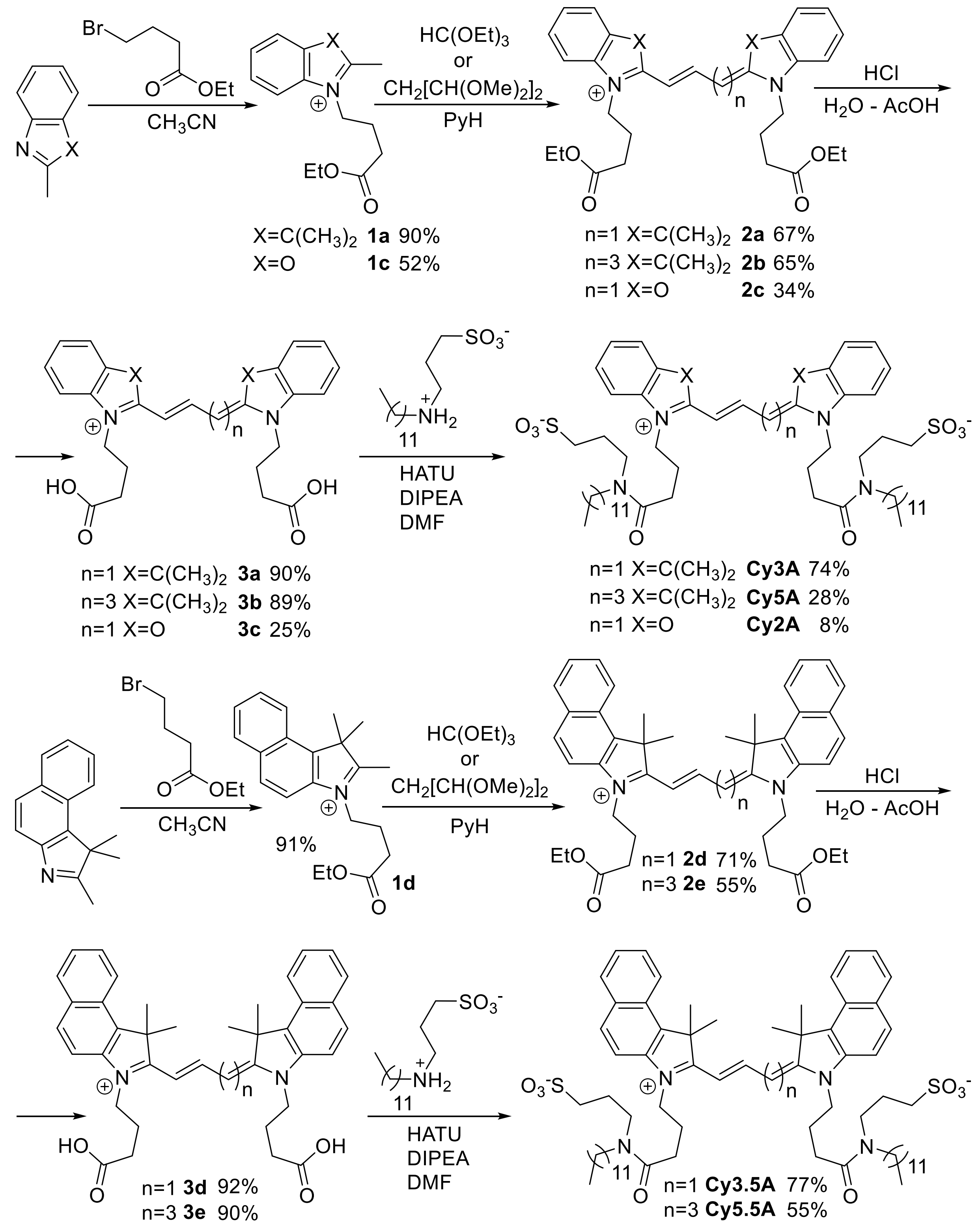


**Scheme S1.** Synthesis of anionic cyanine probes.

**3-(4-ethoxy-4-oxobutyl)-2-methylbenzo[d]oxazol-3-ium** (**1c**)

14.65 g (2.5 eq., 10.74 mL) of ethyl 4-bromobutyrate were added to the saturated solution of KI in 20 mL of dry acetone. The mixture was stirred for 1h, after that the precipitate was filtered off and the solvent was evaporated *in vacuo*.

After this, a solution of 4g (1 eq.) of 2-methylbenzoxazole in 30 mL of dry $CH_3CN$ was added and the resulting mixture was refluxed for 8 days (control by TLC).

After the reaction the inorganic salts were filtered off and the filtrate was diluted with 100 mL of $Et_2O$. The formed mixture was left at 4 °C for 48h, then the formed crystals were filtered off, washed with *ca.* 5 mL of THF to separate the coloured admixtures and refluxed for 10 minutes in 50 mL of THF. The hot supernatant solution was discarded before cooling and the remaining oil was air-dried.

Compound **1c**: yield 5.66 g (52 %) as yellowish solid. **$^1$H NMR (400 MHz, $CDCl_3$) δ ppm** 8.18 - 8.21 (m, 1 H) 7.76 - 7.83 (m, 1 H) 7.64 - 7.71 (m, 2 H) 4.85 (t, *J*=8.30 Hz, 2 H) 4.00 (q, *J*=7.13 Hz, 2 H) 3.40 (s, 3 H) 2.64 (t, *J*=7.00 Hz, 2 H) 2.26 - 2.35 (m, 2 H) 1.16 (t, *J*=7.13 Hz, 3 H). **$^{13}$C NMR (101 MHz, Methanol-*d*$_4$) δ ppm** 172.72 ($C_{carboxyl}$) 167.88 ($C_{ar}$) 147.95 ($C_{ar}$) 129.81 ($C_{ar}$) 129.31 ($C_{ar}$) 128.53 ($C_{ar}$) 115.09 ($C_{ar}$) 113.19 ($C_{ar}$) 60.97 ($C_{al}$) 47.68 ($C_{al}$) 30.50 ($C_{al}$) 23.25 ($C_{al}$) 16.45 ($CH_3$) 14.11 ($CH_3$). **HRMS (ESI), *m/z*:** $[M]^+$ calcd for $C_{14}H_{18}NO_3^+$, 248.1281; found, 248.1282.

**Cyanine 3.5 or 5.5 diethyl ester** (**2d, 2e**) general procedure

4 g of 3-(4-ethoxy-4-oxobutyl)-1,1,2-trimethyl-1H-benzo[e]indol-3-ium were dissolved in 130 mL of dry pyridine upon heating, then the mixture was heated to 110 °C and 1.5 eq of thiethylorthoformate (in case of Cy3.5OEt) or 1.5 eq. of 1,1,3,3-tetramethoxypropane (in case of Cy5.5OEt) were quickly added dropwise to the boiling solution. The mixture was refluxed for 2h (control by TLC).

After the reaction the solvent was evaporated *in vacuo*, then the solid residues were dissolved in DCM and washed with 1M HCl (3 times) and brine (once), dried over $Na_2SO_4$, then the solvent was evaporated *in vacuo*. The crude product was purified by gradient column chromatography (DCM:MeOH 95:5 to 9:1) (performed 2 times for each compound).

Compound **2d**: yield 2.68 g (71 %) as dark violet solid. **$^1$H NMR (400 MHz, Methanol-*d*$_4$) δ ppm** 8.82 (t, *J*=13.5 Hz, 1 H) 8.32 (d, *J*=7.8 Hz, 2 H) 8.09 (d, *J*=8.8 Hz, 2 H) 8.05 (d, *J*=8.0 Hz, 2 H) 7.66 - 7.80 (m, 4 H) 7.50 - 7.62 (m, 2 H) 6.58 (d, *J*=13.5 Hz, 2 H) 4.36 (t, *J*=7.8 Hz, 4 H) 4.15 (q, *J*=7.2 Hz, 4 H) 2.65 (t, *J*=6.8 Hz, 4 H) 2.17 - 2.26 (m, 4 H) 2.14 (s, 12 H) 1.25 (t, *J*=7.2, 1.00 Hz, 6 H).

Compound **2e**: yield 1.8 g (55 %) as dark blue/green solid. **$^1$H NMR (400 MHz, Methanol-*d*$_4$) δ ppm** 8.43 (t, *J*=13.1 Hz, 2 H) 8.27 (d, *J*=8.5 Hz, 2 H) 8.05 (d, *J*=9.0 Hz, 2 H) 8.02 (d, *J*=8.3 Hz, 2 H) 7.64 - 7.71 (m, 4 H) 7.48 - 7.55 (m, 2 H) 6.69 (t, *J*=12.5 Hz, 1 H) 6.45 (d, *J*=13.8 Hz, 2 H) 4.30 (br t, *J*=7.8 Hz, 4 H) 4.19 (q, *J*=7.0 Hz, 4 H) 2.63 (t, *J*=6.7 Hz, 4 H) 2.11 - 2.24 (m, 4 H) 2.05 (s, 12 H) 1.28 (t, *J*=7.0 Hz, 6 H). **$^{13}$C NMR (101 MHz, $CDCl_3$) δ ppm** 174.19 ($C_{carboxyl}$) 172.96 ($C_{ar}$) 153.10 ($C_{ar}$) 139.45 ($C_{ar}$) 133.83 ($C_{ar}$) 131.78 ($C_{ar}$) 130.61 ($C_{ar}$) 129.97 ($C_{ar}$) 128.18

($C_{ar}$) 127.66 ($C_{ar}$) 124.94 ($C_{ar}$) 122.26 ($C_{ar}$) 110.67 ($C_{ar}$) 103.53 ($C_{ar}$) 60.78 ($C_{al}$) 51.18 ($C_{al}$) 43.57 ($C_{al}$) 30.59 ($C_{al}$) 27.68 ($C_{al}$) 22.47 ($C_{al}$) 14.22 ($CH_3$). **HRMS (ESI), *m/z*:** $[M]^+$ calcd for $C_{45}H_{51}N_2O_4^+$, 683.3843; found, 683.3850.

**Cyanine 2 diethyl ester** (**2c**)

5 g of 3-(4-ethoxy-4-oxobutyl)-2-methylbenzo[d]oxazol-3-ium were dissolved in 100 mL of dry pyridine upon heating, then the mixture was heated to 110 °C and 2.50 g (2.81 mL, 1.3 eq.) of thiethylortoformate were quickly added dropwise to the boiling solution. The mixture was refluxed for 5h (control by TLC).

After the reaction the solvent was evaporated *in vacuo*, then the solid residues were dissolved in DCM and washed with 1M HCl (3 times) and brine (once), dried over $Na_2SO_4$, then the solvent was evaporated *in vacuo*.

The crude product was purified by gradient column chromatography (DCM:MeOH 95:5 to 85:15) and consecutive second column chromatography (DCM:MeOH 95:5).

Compound **2c**: yield 1.34 g (34 %) as dark red solid. **$^1$H NMR (400 MHz, Methanol-*d*$_4$) δ ppm** 8.44 (t, *J*=13.2 Hz, 1 H) 7.44 - 7.49 (m, 4 H) 7.40 (td, *J*=7.7, 1.0 Hz, 2 H) 7.33 (dq, *J*=7.7, 1.0 Hz, 2 H) 6.48 (t, *J*=13.2 Hz, 2 H) 4.25 (quin, *J*=7.8 Hz, 4 H) 4.14 (q, *J*=7.3 Hz, 4 H) 2.63 (t, *J*=6.4 Hz, 4 H) 2.10 - 2.25 (m, 4 H) 1.27 (t, *J*=7.3 Hz, 6 H). **$^{13}$C NMR (101 MHz, Methanol-*d*$_4$) δ ppm** 173.15 ($C_{carboxyl}$) 161.92 ($C_{ar}$) 148.03 ($C_{ar}$) 146.92 ($C_{ar}$) 131.30 ($C_{ar}$) 126.15 ($C_{ar}$) 125.04 ($C_{ar}$) 110.68 ($C_{ar}$) 110.09 ($C_{methylenic}$) 86.85 ($C_{methylenic}$) 60.72 ($C_{al}$) 43.45 ($C_{al}$) 30.19 ($C_{al}$) 22.79 ($C_{al}$) 14.19 ($CH_3$). **HRMS (ESI), *m/z*:** $[M]^+$ calcd for $C_{29}H_{33}N_2O_6^+$, 505.2333; found, 505.2348.

**Cyanine diacid** (**3a-e**) general procedure.

900 mg of corresponding cyanine diethyl ester were dissolved in 6 ml of glacial acetic acid, then 4 ml of $H_2O$ and 2 ml of $HCl_{conc}$ were added. The reaction mixture was refluxed for 40 minutes (control by TLC).

After the reaction the solvents were evaporated *in vacuo*, and the solid residue was washed twice with acidified (pH 4) water on filter, then air-dried. In case of compound **3c** an additional purification by gradient column chromatography (DCM:MeOH:HCOOH 98:2:2 to 85:15:3) was needed.

Compound **3a**: yield 739 mg (90 %) as dark red solid. **$^1$H NMR (400 MHz, DMSO-*d*$_6$) δ ppm** 8.36 (t, *J*=13.3 Hz, 1 H) 7.65 (d, *J*=7.3 Hz, 2 H) 7.41 - 7.53 (m, 4 H) 7.30 (t, *J*=7.4 Hz, 2 H) 6.54 (d, *J*=13.3 Hz, 2 H) 4.15 (t, *J*=7.2 Hz, 4 H) 2.43 (t, *J*=7.2 Hz, 4 H) 1.90 - 2.03 (m, 4 H) 1.71 (s, 12 H).

Compound **3b**: yield 734 mg (89 %) as dark blue solid. **$^1$H NMR (400 MHz, Methanol-*d*$_4$) δ ppm** 8.28 (t, *J*=13.2 Hz, 2 H) 7.49 (d, *J*=7.3 Hz, 2 H) 7.33 - 7.45 (m, 4 H) 7.23 - 7.31 (m, 2 H) 6.63 (t, *J*=12.4 Hz, 1 H) 6.37 (d, *J*=13.8 Hz, 2 H) 4.11 - 4.20 (m, 4 H) 2.48 - 2.57 (m, 4 H) 2.00 - 2.11 (m, 4 H) 1.73 (s, 12 H).

Compound **3c**: yield 205 mg (25 %) as dark yellow solid. **$^1$H NMR (400 MHz, DMSO-*d*$_6$) δ ppm** 11.79 (br s, 2 H) 8.16 - 8.37 (m, 1 H) 7.33 - 7.89 (m, 1 H) 6.73 - 7.28 (m, 7 H) 6.30 -

6.73 (m, 2 H) 3.92 - 4.09 (m, 2 H) 3.51 - 3.75 (m, 2 H) 2.11 - 2.36 (m, 4 H) 1.44 - 1.70 (m, 4 H). **$^{13}$C NMR (101 MHz, Methanol-*d*$_4$) δ ppm** 174.66 (COOH) 173.06 ($C_{ar}$) 154.11 ($C_{ar}$) 153.85 ($C_{ar}$) 130.37 ($C_{ar}$) 129.58 ($C_{ar}$) 120.01 ($C_{ar}$) 117.58 ($C_{ar}$) 117.10 ($C_{ar}$) 109.33 ($C_{ar}$) 60.13 ($C_{al}$) 31.15 ($C_{al}$) 23.28 ($C_{al}$). **HRMS (ESI), *m/z*:** $[M-2H+2Na]^+$ calcd for $C_{25}H_{23}N_2Na_2O_6^+$, 493.1346; found, 493.1366.

Compound **3d**: yield 765 mg (92 %) as dark violet solid. **$^1$H NMR (400 MHz, DMSO-*d*$_6$) δ ppm** 8.56 (t, *J*=13.3 Hz, 1 H) 8.29 (d, *J*=9.5 Hz, 2 H) 8.11 (d, *J*=9.0 Hz, 2 H) 8.08 (d, *J*=8.3 Hz, 2 H) 7.83 (d, *J*=8.8 Hz, 2 H) 7.68 (t, *J*=7.7 Hz, 2 H) 7.53 (t, *J*=7.4 Hz, 2 H) 6.65 (d, *J*=13.3 Hz, 2 H) 4.27 (br t, *J*=6.8 Hz, 4 H) 2.37 (br t, *J*=6.27 Hz, 4 H) 2.05 - 2.10 (m, 2 H) 2.01 (s, 12 H) 1.96 - 1.99 (m, 2 H). **$^{13}$C NMR (101 MHz, Methanol-*d*$_4$) δ ppm** 175.57 (COOH) 174.74 ($C_{ar}$) 148.78 ($C_{ar}$) 140.04 ($C_{ar}$) 133.57 ($C_{ar}$) 131.96 ($C_{ar}$) 130.97 ($C_{ar}$) 130.43 ($C_{ar}$) 128.33 ($C_{ar}$) 127.94 ($C_{ar}$) 125.46 ($C_{ar}$) 122.63 ($C_{ar}$) 112.08 ($C_{ar}$) 102.98 ($C_{ar}$) 51.04 ($C_{al}$) 44.06 ($C_{al}$) 32.26 ($C_{al}$) 27.58 ($C_{al}$) 23.72 ($CH_3$). **HRMS (ESI), *m/z*:** $[M]^+$ calcd for $C_{39}H_{41}N_2O_4^+$, 601.3061; found, 601.3079.

Compound **3e**: yield 751 mg (90 %) as dark blue/green solid. **$^1$H NMR (400 MHz, DMSO-*d*$_6$) δ ppm** 12.25 (br s, 2 H) 8.50 (t, *J*=13.2 Hz, 2 H) 8.27 (d, *J*=8.5 Hz, 2 H) 8.10 (d, *J*=8.8 Hz, 2 H) 8.07 (d, *J*=8.5 Hz, 2 H) 7.78 (d, *J*=9.0 Hz, 2 H) 7.69 (t, *J*=7.8 Hz, 2 H) 7.53 (t, *J*=7.7 Hz, 2 H) 6.61 (t, *J*=12.5 Hz, 1 H) 6.44 (d, *J*=13.6 Hz, 2 H) 4.25 (br t, *J*=7.2 Hz, 4 H) 2.22 - 2.36 (m, 2 H) 2.03 - 2.16 (m, 4 H) 1.98 (s, 12 H) 1.90 - 1.96 (m, 2 H). **$^{13}$C NMR (101 MHz, Methanol-*d*$_4$) δ ppm** 174.36 (COOH) 174.29 ($C_{ar}$) 153.54 ($C_{ar}$) 140.15 ($C_{ar}$) 133.64 ($C_{ar}$) 131.79 ($C_{ar}$) 130.78 ($C_{ar}$) 130.40 ($C_{ar}$) 128.20 ($C_{ar}$) 128.10 ($C_{ar}$) 126.11 ($C_{ar}$) 125.26 ($C_{ar}$) 122.62 ($C_{ar}$) 111.93 ($C_{ar}$) 103.37 ($C_{ar}$) 51.24 ($C_{al}$) 43.42 ($C_{al}$) 30.84 ($C_{al}$) 27.25 ($C_{al}$) 22.88 ($CH_3$). **HRMS (ESI), *m/z*:** $[M]^+$ calcd for $C_{41}H_{43}N_2O_4^+$, 627.3217; found, 627.3235.

**Cyanine dialkyl disulfonate** (**Cy3A, Cy5A, Cy2A, Cy3.5A, Cy5.5A**) general procedure

30 mg of the corresponding cyanine diacid were mixed with 2.1 equiv. of HATU and 3 equiv. of DIPEA in 1 mL of dry DMF. After 5 minutes, a solution of 2.1 equiv. of 3-(dodecylammonio)propane-1-sulfonate together with 3 equiv. of DIPEA in dry DMF was added and the reaction mixture was stirred for 24h at r.t. (control by TLC) under Ar atmosphere.

After the reaction the solvent was evaporated *in vacuo*, and the crude product was purified by exclusive column chromatography (LH 20, DCM:MeOH 1:1 for all compounds) and, consecutively, by preparative TLC ($SiO_2$, DCM:MeOH 9:1 for **Cy3A**, **Cy5A**, **Cy3.5A**; DCM:MeOH 85:15 for **Cy5.5A**; DCM:MeOH 82:18 for **Cy2A**).

Compound **Cy3A**: yield 41 mg (74 %) as dark red solid. **$^1$H NMR (400 MHz, DMSO-*d*$_6$) δ ppm** 8.37 (t, *J*=13.3 Hz, 1 H) 7.58 - 7.70 (m, 2 H) 7.38 - 7.58 (m, 4 H) 7.23 - 7.34 (m, 2 H) 6.48 - 6.85 (m, 2 H) 4.09 - 4.27 (m, 4 H) 3.08 - 3.29 (m, 8 H) 2.54 - 2.63 (m, 4 H) 2.28 - 2.46 (m, 4 H) 1.88-2.02 (m, 4 H) 1.77 - 1.87 (m, 4 H) 1.70 (br s, 12 H) 1.34-1.46 (m, 4 H) 1.14-1.34 (m, 36 H) 0.80-0.89 (m, 6 H). $^{13}$C NMR spectrum could not be obtained due to probe aggregation. **HRMS (ESI), *m/z*:** $[M]^-$ calcd for $C_{61}H_{97}N_4O_8S_2^-$, 1077.6753; found, 1077.6771.

Compound **Cy5A**: yield 15 mg (28 %) as dark blue solid. **$^1$H NMR (400 MHz, DMSO-*d*$_6$) δ ppm** 8.34 (t, *J*=13.1 Hz, 2 H) 7.61 (dd, *J*=7.3, 4.0 Hz, 2 H) 7.35 - 7.52 (m, 4 H) 7.19-7.28 (m, 2 H) 6.29 - 6.67 (m, 3 H) 4.07-4.20 (m, 4 H) 3.20 - 3.32 (m, 4 H) 3.17 (d, *J*=5.3 Hz, 4 H) 2.53 - 2.59 (m, 2 H) 2.38 - 2.49 (m, 6 H) 1.85 - 1.99 (m, 4 H) 1.76 - 1.85 (m, 4 H) 1.69 (s, 12 H) 1.35 - 1.47 (m, 4 H) 1.14 - 1.28 (m, 36 H) 0.79 - 0.87 (m, 6 H). $^{13}$C NMR spectrum could not be obtained due to probe aggregation. **HRMS (ESI), *m/z*:** [M]$^-$ calcd for $C_{63}H_{99}N_4O_8S_2^-$, 1103.6910; found, 1103.6912.

Compound **Cy2A**: yield 8 mg (8 %) as dark yellow solid. **$^1$H NMR (400 MHz, DMSO-*d*$_6$) δ ppm** 8.31 (td, *J*=13.4, 5.3 Hz, 1 H) 7.73 (m, 4 H) 7.36 - 7.54 (m, 4 H) 6.05 - 6.32 (m, 2 H) 4.07 - 4.32 (m, 4 H) 3.11 - 3.28 (m, 8 H) 2.56 - 2.69 (m, 4 H) 2.31 - 2.45 (m, 4 H) 1.94 - 2.07 (m, 4 H) 1.67 - 1.86 (m, 4 H) 1.33 - 1.47 (m, 4 H) 1.10 - 1.32 (m, 36 H) 0.85 (m, 6 H). $^{13}$C NMR spectrum could not be obtained due to probe aggregation. **HRMS (ESI), *m/z*:** [M]$^-$ calcd for $C_{55}H_{85}N_4O_{10}S_2^-$, 1025.5713; found, 1025.5754.

Compound **Cy3.5A**: yield 40 mg (77 %) as dark violet solid. **$^1$H NMR (400 MHz, Methanol-*d*$_4$) δ ppm** 8.59 (t, *J*=13.5 Hz, 1 H) 8.28 (d, *J*=8.5 Hz, 2 H) 8.12 (t, *J*=9.0 Hz, 2 H) 8.07 (d, *J*=8.3 Hz, 2 H) 7.81 - 7.89 (m, 2 H) 7.67 (t, *J*=7.8 Hz, 2 H) 7.52 (t, *J*=7.5 Hz, 2 H) 6.57 - 6.86 (m, 2 H) 4.23 - 4.39 (m, 4 H) 3.40 - 3.51 (m, 2 H) 3.23 - 3.29 (m, 2 H) 3.10 - 3.23 (m, 4 H) 2.53 - 2.69 (m, 4 H) 2.32 - 2.47 (m, 4 H) 2.02 (s, 12 H) 1.79 - 1.95 (m, 4 H) 1.47 - 1.79 (m, 4 H) 1.30 - 1.40 (m, 4 H) 1.16 - 1.27 (m, 36 H) 0.78 - 0.87 (m, 6 H). $^{13}$C NMR spectrum could not be obtained due to probe aggregation. **HRMS (ESI), *m/z*:** [M]$^-$ calcd for $C_{69}H_{101}N_4O_8S_2^-$, 1177.7066; found, 1177.7069.

Compound **Cy5.5A**: yield 28 mg (55 %) as dark blue/green solid. **$^1$H NMR (400 MHz, DMSO-*d*$_6$) δ ppm** 8.46 (t, *J*=13.1 Hz, 2 H) 8.25 (d, *J*=8.5 Hz, 2 H) 8.02 - 8.13 (m, 4 H) 7.83 (t, *J*=8.3 Hz, 1 H) 7.77 (t, *J*=9.0 Hz, 1 H) 7.67 (t, *J*=7.0 Hz, 2 H) 7.51 (t, *J*=6.8 Hz, 2 H) 6.60 - 6.75 (m, 1 H) 6.37 - 6.58 (m, 2 H) 4.19 - 4.34 (m, 4 H) 3.40 - 3.55 (m, 2 H) 3.12 - 3.27 (m, 6 H) 2.56 - 2.69 (m, 2 H) 2.31 - 2.48 (m, 6 H) 2.03 - 2.07 (m, 2 H) 1.97 (d, *J*=3.3 Hz, 12 H) 1.64 - 1.90 (m, 6 H) 1.32 - 1.45 (m, 4 H) 1.17 (m, 36 H) 0.74 - 0.89 (m, 6 H). $^{13}$C NMR spectrum could not be obtained due to probe aggregation. **HRMS (ESI), *m/z*:** [M]$^-$ calcd for $C_{71}H_{103}N_4O_8S_2^-$, 1203.7223; found, 1203.7243.

**Preparation of liposomes.** All types of LUVs used were prepared by the following procedure. A stock solution of corresponding lipid(s) in chloroform was placed into a round-neck flask, after which the solvent was evaporated *in vacuo* and phosphate buffer (20 mM, pH 7.4) was added. After all the solid was dissolved a suspension of multilamellar vesicles was extruded by using a Lipex Biomembranes extruder (Vancouver, Canada). The size of the filters was first 0.2 μm (7 passages) and thereafter 0.1 μm (10 passages). This generates monodisperse LUVs with a mean diameter of 0.12 μm as measured with a Malvern Zetamaster 300 (Malvern, U.K.).

**Cell Lines, Culture Conditions, and Treatment.** KB (ATCC CCL-17) and MDA-MB-231 (ATCC HTB-26) cells were grown in Dulbecco`s Modified Eagle Medium (DMEM, Gibco

Invitrogen), supplemented with 10% fetal bovine serum (FBS, Lonza), 1% l-Glutamine (Sigma Aldrich) and 1% antibiotic solution (penicillin-streptomycin, Gibco-Invitrogen) at 37 °C in a humidified 5% $CO_2$ atmosphere. Cells were seeded onto a chambered coverglass (IBIDI) at a density of $7\times10^4$ cells/well 24 h before the microscopy measurement. For cellular microscopy experiments, the attached KB cells in IBIDI dishes were washed twice with warm Hank`s balanced salt solution (HBSS, Gibco Invitrogen), after that 1 mL of dye solution in HBSS was added and the cells were incubated for 10 min. at r.t.

**Fluorescence microscopy.** Imaging of KB cells was performed using Nikon Ti-E inverted microscope, equipped with CFI Plan Apo ×60 oil (NA = 1.4) objective, X-Light spinning disk module (CREST Optics) and a Hamamatsu Orca Flash 4 sCMOS camera with a bandpass filter $531 \pm 40$ nm (Semrock) or $593 \pm 40$ nm (Semrock) or longpass filter 647 nm (Semrock). The excitation in confocal mode was provided by a 488 nm or 532 nm or 638 nm diode laser (OXXIUS). The exposure time in confocal mode was set to 500 ms per image frame. All the images were recorded using NIS Elements and then processed using Fiji software.

Fluorescence confocal microscopy of MDA-MB-231 were performed on a Leica TCS SPE-II microscope with a HC PL APO CS2 63x/1.40 OIL objective. The excitation of Hoechst 33342 was performed with a 405nm 10 mW laser and the emission was detected around 450-500 nm. The excitation of Cy2A or Cy3A/Cy3.5A or Cy5A/Cy5.5A in confocal mode was provided by a 488 nm or 532 nm or 638 nm diode laser respectively. For colocalization experiments, 500 µL of CyXA and WGA-Alexa488 (Wheat Germ Agglutinin (WGA), Thermofischer) of 100 nM each dissolved in HBSS solution and added to live cells to reach final concentration of 50nM. 25 nM of CyXA and MBCyX probes were used for brightness and internalization comparison. Cells were incubated at room temperature for 10 or 15 min to perform the imaging.

**Intracranial dye injection and cranial window implantation.** All animal experiments were conducted in accordance with institutional guidelines and approved by the Government of Upper Bavaria. 8-week old C56/Bl6 mice were obtained from Charles River Laboratories (Kisslegg, Germany). Cranial window implantation was performed as decribed in [7]. Before use, surgical tools were sterilized in a glass-bead sterilizer (FST). Mice were anesthetized by an intra-peritoneal injection of MMF (medetomidine (0.5 mg/kg), fentanyl (0.05 mg/kg), and midazolam (5 mg/kg)). Subsequently, mice were placed onto a heating blanket (37 °C) and the head was fixed in a stereotactic frame. Eyes were protected from drying by applying eye ointment (Bepanthen, Bayer). The scalp was washed with swabs soaked with 70 % ethanol. A flap of skin covering the cranium was excised using small scissors. The periosteum was scraped away with a scalpel. The prospective craniotomy location (1.5-2.5 mm AP and 0-4 mm ML relative to bregma) was marked with a biopsy punch (diameter 4 mm, Integra LifeSciences). The exposed skull around the area of interest was covered with a thin layer of dental acrylic (iBond Self Etch, Hereaus Kulzer) and hardened with a LED polymerization lamp (Demi Plus, Kerr). A dental drill

(Schick Technikmaster C1, Pluradent) was used to thin the skull around the marked area. After applying a drop of sterile phosphate buffered saline (DPBS, Gibco, Life Technologies) on the craniotomy the detached circular bone flap was removed with forceps. For *in vivo* imaging, a concentrated stock solution of Cy3.5A in DMSO was added to the 1 mM solution of Me-β-Cyclodextrin in PBS to create a 20 µM solution of Cy3.5A, immediately used for imaging. 300 nl of probe solution were injected at 200 µm depth from the brain surface via a Nanoliter 2000 Injector (World Precision Instruments) at a speed of 30 nl/min. A circular coverslip (4 mm diameter, VWR International) was placed onto the craniotomy and glued to the skull with histoacryl adhesive (Aesculap). The exposed skull was covered with dental acrylic (Tetric Evoflow A1 Fill, Ivoclar Vivadent) and a head-post was attached parallel to the window for head-fixing mice subsequently under the 2-photon laser scanning microscope (LSM 7 MP, Carl Zeiss Ltd.).

***In vivo* 2-photon microscopy.** *In vivo* 2-photon imaging was performed using a multiphoton LSM 7 MP microscope (Zeiss) equipped with a Ti:Sa laser (Chameleon Vision II from Coherent, Glasgow, Scotland), a 20x water immersion objective (W Plan-Apochromat 20x/1.0 NA, Zeiss) and a motorized stage as decribed in [8]. The membrane probe was excited at 830 nm and the emission was collected after a LP>750 nm filter by a non-descanned detector (photomultiplier tube GaAsP, Zeiss). Throughout the imaging session mice were kept on a heating pad to keep body temperature at 37 °C (Fine Science Tools GmbH). For overview images, 3D stacks of 100 µm depth with 3 µm axial resolution and 1024×1024 pixels per image frame (0.4 µm/pixel) were acquired. To resolve dendritic spines and axonal boutons, high-resolution images from single neurites were taken with 0.7 µm axial resolution and 512 × 256 pixels per image frame (0.1 µm per pixel) with the laser power kept below 50 mW to avoid phototoxicity. Fig. 5 A,B was created with BioRender.com.

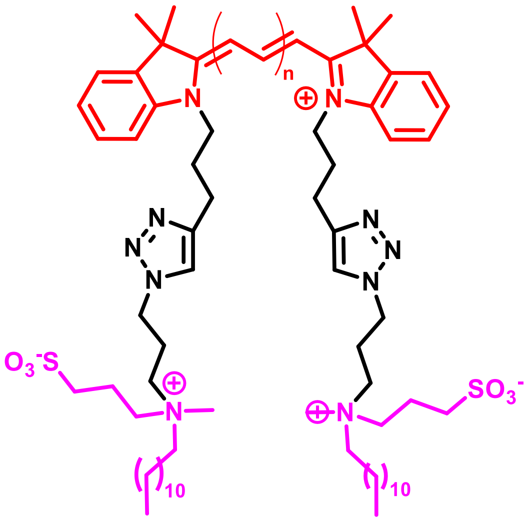

**Membright-Cy3; n = 1**
**Membright-Cy3.5; n = 1**
**Membright-Cy5; n = 2**
**Membright-Cy5.5; n = 2**

**Fig. S1.** Chemical structure of MemBright probes: parent analogues of the membrane probes of the present study.

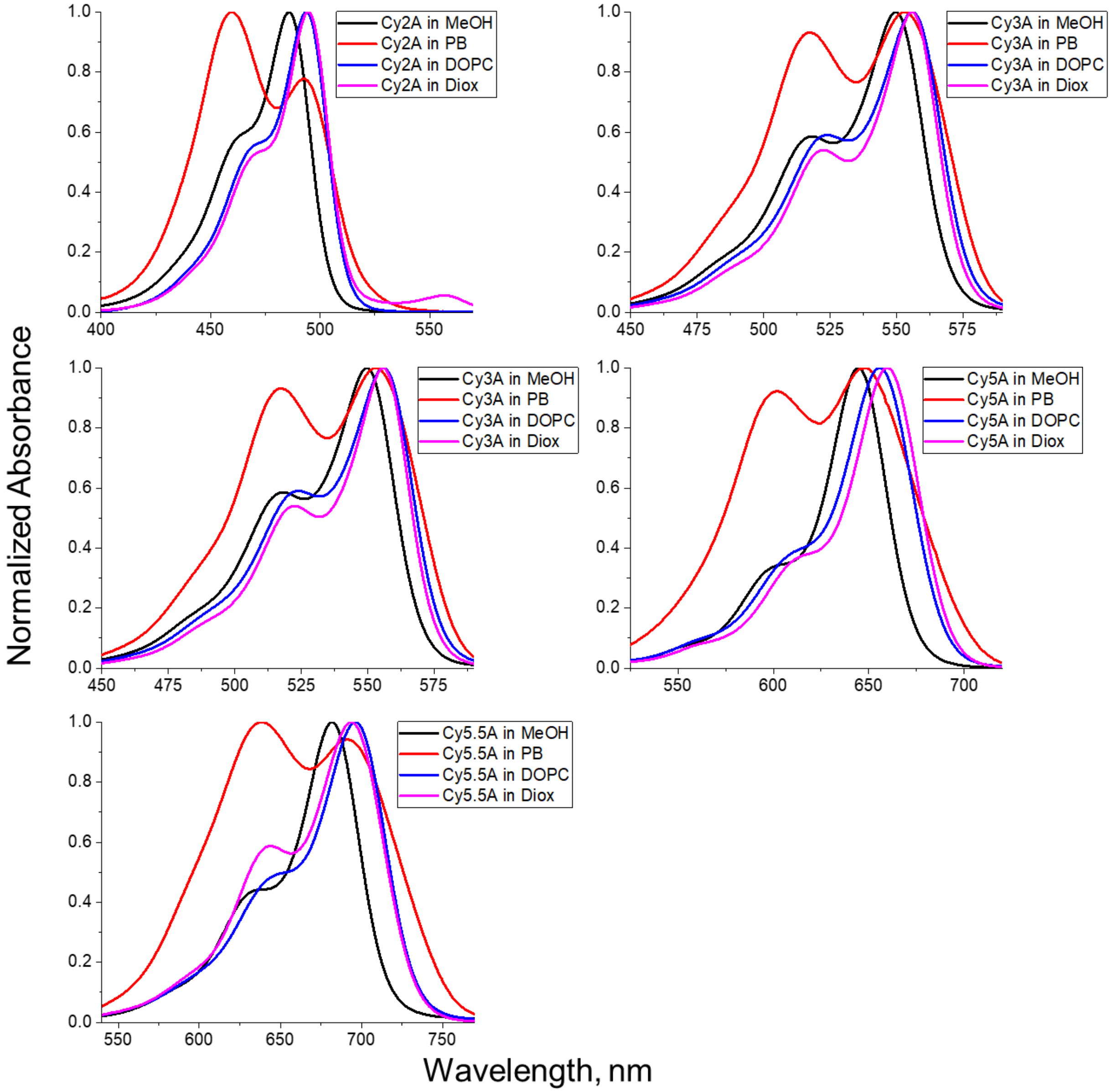


**Fig. S2.** Normalized absorption spectra of anionic cyanine probes in solvents and lipid vesicles. Probe concentration was 1 µM for Cy2A and Cy3A and 0.5 µM for Cy3.5A, Cy5A and Cy5.5A. Total lipid concentration was 400 µM.

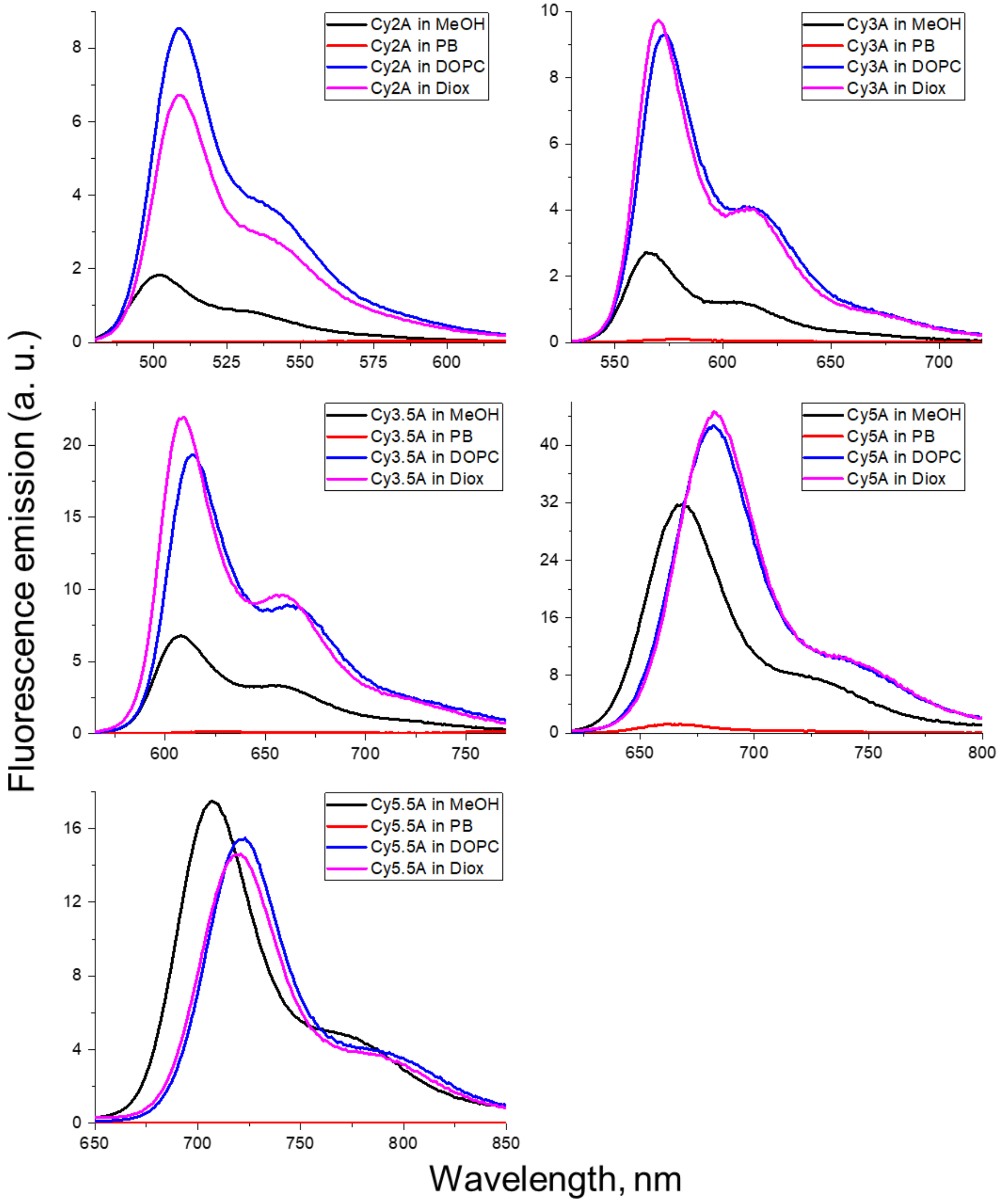


**Fig. S3.** Normalized emission spectra of anionic cyanine probes in solvents and lipid vesicles. All the spectra were corrected by dividing by the absorption on the excitation wavelength. Probe concentration was 1 µM for Cy2A and Cy3 and 0.5 µM for Cy3.5, Cy5 and Cy5.5. Total lipid concentration was 400 µM. Excitation wavelength was 470 nm for Cy2A, 540 nm for Cy3A, 546 nm for Cy3.5A, 601 nm for Cy5A and 623 nm for Cy5.5A.

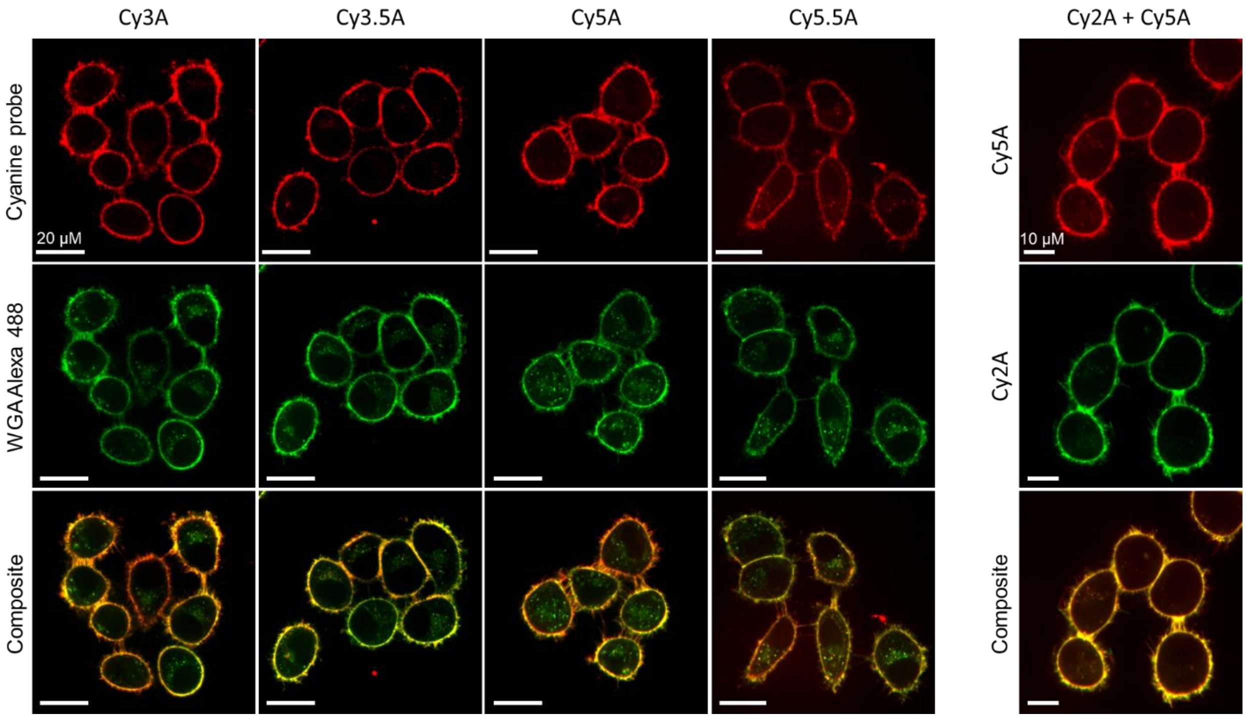


**Fig. S4.** Colocalization of anionic cyanine probes with commercial WGA-Alexa 488 conjugate in live KB cells. Concentrations of all dyes used were 50 nM. Scale bar: 20 µM for Cy3A, Cy3.5A, Cy5A and Cy5.5A panels; 10 µM for Cy2A+Cy5A panel.

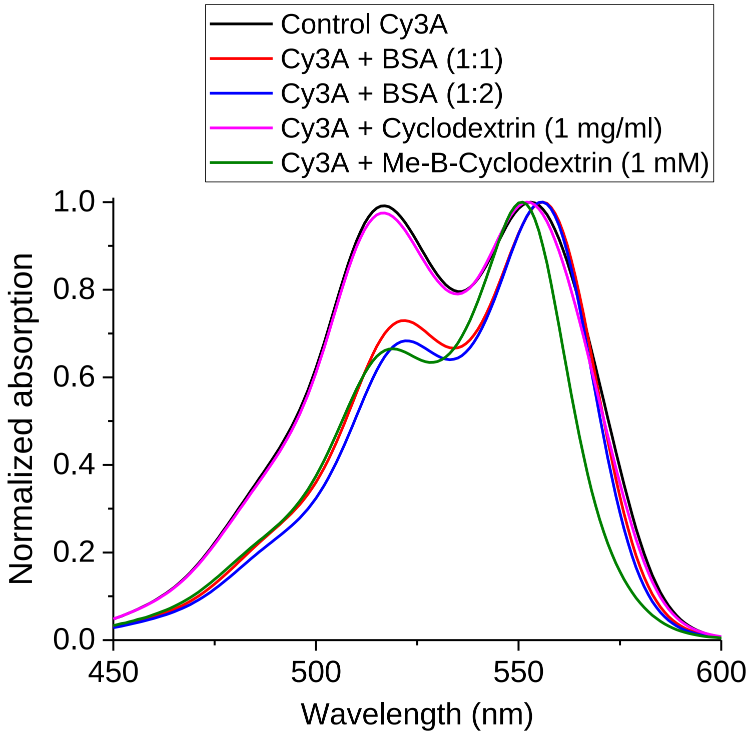


**Fig. S5.** Normalized absorption spectra of Cy3A (20 μM) in PBS with or without various delivery agents.

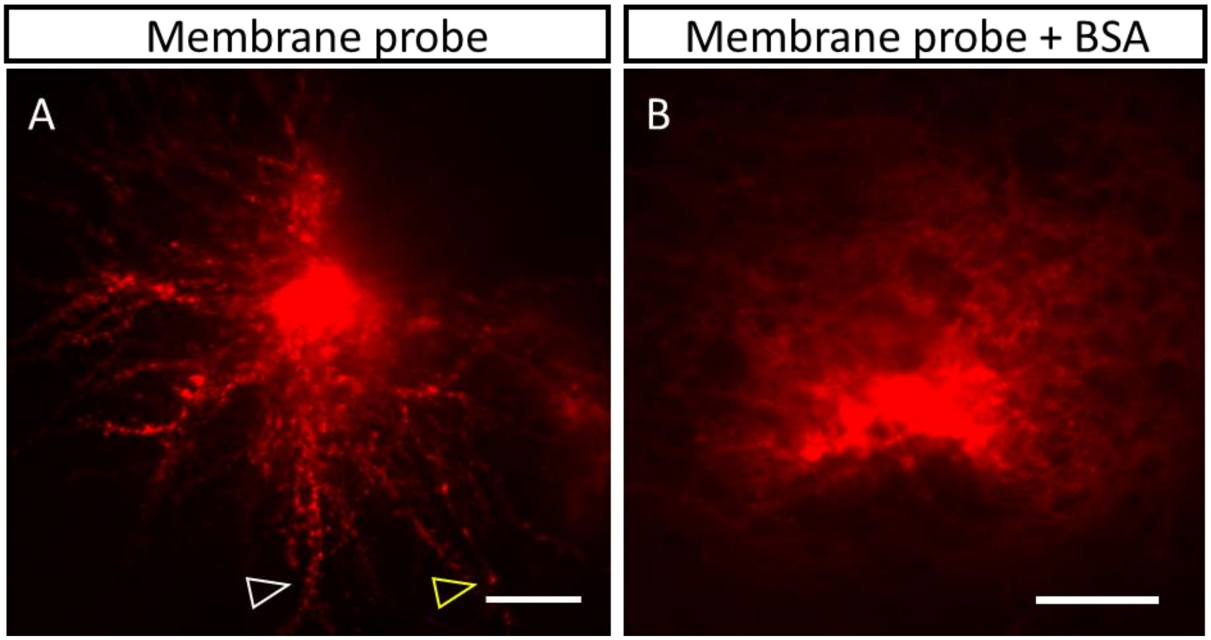


**Fig. S6.** *In vivo* 2-photon microscopy of neurons stained by different probe formulations injected into the mouse brain parenchyma with experiment design presented at Fig.5A,B. A. Representative *in vivo* 2-photon maximum intensity projection image of pure membrane probe injected into the mouse brain without any additive. Sparse staining of dendrite (white arrow) and axon (yellow arrow). B. Representative *in vivo* 2-photon image of membrane probe formulated in Bovine Serum Albumin solution (BSA) injected into the mouse brain. Nonspecific staining. Scale bar – 20 µm.
